# Binding of a Macrocyclic Nonapeptide Inhibitor to Histone Deacetylase 6

**DOI:** 10.64898/2026.10.10.758213

**Authors:** Abraham Eafa, Parisa Hosseinzadeh, David Baker, David W. Christianson

**Affiliations:** Roy and Diana Vagelos Laboratories, Department of Chemistry, University of Pennsylvania, Philadelphia, Pennsylvania, 19104-6323, United States; Department of Bioengineering, Knight Campus, University of Oregon, Eugene, OR 97403 United States; Department of Biochemistry, Institute for Protein Design, University of Washington, Seattle, WA 98195 United States

**Author notes:** Corresponding author: **David W. Christianson –** Roy and Diana Vagelos Laboratories, Department of Chemistry, University of Pennsylvania, Philadelphia, Pennsylvania 19104-6323, United States.

## Abstract

Histone deacetylase 6 (HDAC6) plays an essential role in the regulation of post-translational lysine acetylation. Myriad biological processes depend on the precise regulation of lysine acetylation-deacetylation cycles, and their dysregulation is linked to human pathologies such as cancer and neurodegeneration. Broad-specificity HDAC inhibitors are efficacious in treating disease, but the inhibitors currently approved for clinical use often cause deleterious side effects due to inhibition of multiple HDAC isozymes as well as other zinc enzymes. Cyclic peptides comprise novel platforms that, in principle, can be tailored for affinity and selectivity. A key advantage of cyclic peptide design is the ability to incorporate both canonical and noncanonical amino acids. As part of our program focusing on cyclic peptide HDAC inhibitors, we now report the 1.9-Å resolution crystal structure of HDAC6 CD2 complexed with macrocyclic nonapeptide **1**. The structure reveals zinc coordination by a noncanonical thiolate-containing amino acid, plus direct and water-mediated enzyme-inhibitor hydrogen bonds that contribute to nanomolar inhibitory potency. Structural comparisons with previously reported HDAC6 CD2 complexes with tetrapeptide and octapeptide macrocycles reveal similar networks of direct and water-mediated interactions. Given the surge of interest in cyclic peptide inhibitors, fueled in part by advances in computational design approaches, the crystal structure of the HDAC6 CD2-**1** complex sets an important foundation for further development of macrocyclic peptide inhibitors.

**For Table of Contents Use Only:** 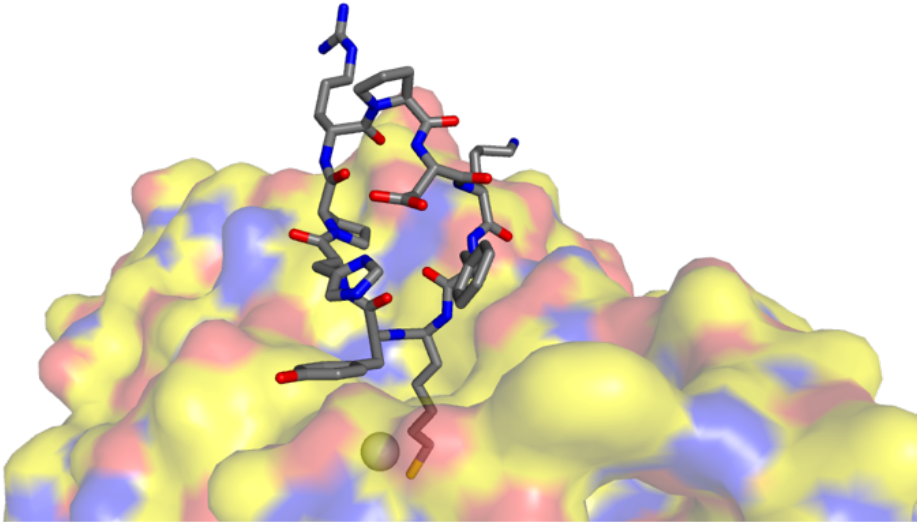

## Introduction

Reversible post-translational modifications (PTMs) of lysine side chains, such as acetylation,^1–4^ succinylation,^5,6^ myristoylation,^7,8^ and more,^9–11^ play an important role in regulating protein function in the cell. PTMs alter the covalent structure (i.e., primary structure) of a protein, and in doing so they can serve as molecular rheostats to modulate biological processes. For example, lysine acetylation serves prominent regulatory functions in epigenetics,^4,12,13^ cell signaling,^14,15^ and metabolism.^16–18^ Lysine acetyltransferases (also known as histone acetyltransferases, HATs) install acetyl groups on target lysine residues using acetyl-CoA as an acetyl donor,^19–21^ and lysine deacetylases (also known as histone deacetylases, HDACs) hydrolyze acetyllysine to yield lysine and acetate.^22,23^ Dysregulation of lysine acetylation/deacetylation is observed in several human pathologies, including cancer and neurodegenerative diseases.^24^ Accordingly, HATs and HDACs are important targets for therapeutic drug design.

Zinc-dependent HDACs are grouped into classes I, IIa, IIb, and IV based on phylogenetic analysis.^25^ Class III HDACs are better known as sirtuins, but these enzymes do not require Zn^2+^ and are structurally and mechanistically distinct.^26^ Of particular interest is the class IIb enzyme HDAC6, which stands out among the greater HDAC family because it contains two catalytic domains, CD1 and CD2, and because it is the largest HDAC (molecular weight = 131 kDa).^27,28^ CD1 exhibits narrow substrate specificity and prefers substrates bearing C-terminal acetyllysine residues,^29,30^ while CD2 exhibits much broader substrate specificity and is responsible for the deacetylation and regulation of cytoplasmic proteins such as *a*-tubulin,^31^ tau,^32^ and cortactin.^33^

The general design of an HDAC inhibitor includes a zinc-binding group, a linker segment, and a capping group,^34,35^ although there are intriguing examples of inhibitors lacking zinc-binding groups.^36–38^ The zinc-binding group coordinates to the catalytic Zn^2+^ ion deep in the active site pocket. The linker segment typically interacts with the walls of the active site, and the capping group interacts with residues at the mouth of the active site. Currently, four HDAC inhibitors are approved for clinical use by the U.S. Food and Drug Administration: Vorinostat, Belinostat, Givinostat, and Romidepsin (Figure 1). All four of these inhibitors are pan-HDAC inhibitors to some degree, in that they are active against multiple zinc-dependent HDACs; moreover, some are active against other enzymes, such as the zinc-dependent metallo beta-lactamase MBLAC2.^39^ Due to their general lack of isozyme selectivity, use of these drugs is usually accompanied by deleterious off-target effects.^40^ Additionally, hydroxamate zinc-binding groups in Vorinostat, Belinostat, and Givinostat can lead to mutagenicity and cytotoxicity through metabolism of the hydroxamic acid.^41^ These disadvantages motivate the continuing search for new inhibitor designs that will serve as frameworks for exploring the molecular determinants of affinity and selectivity.

**Figure 1.**
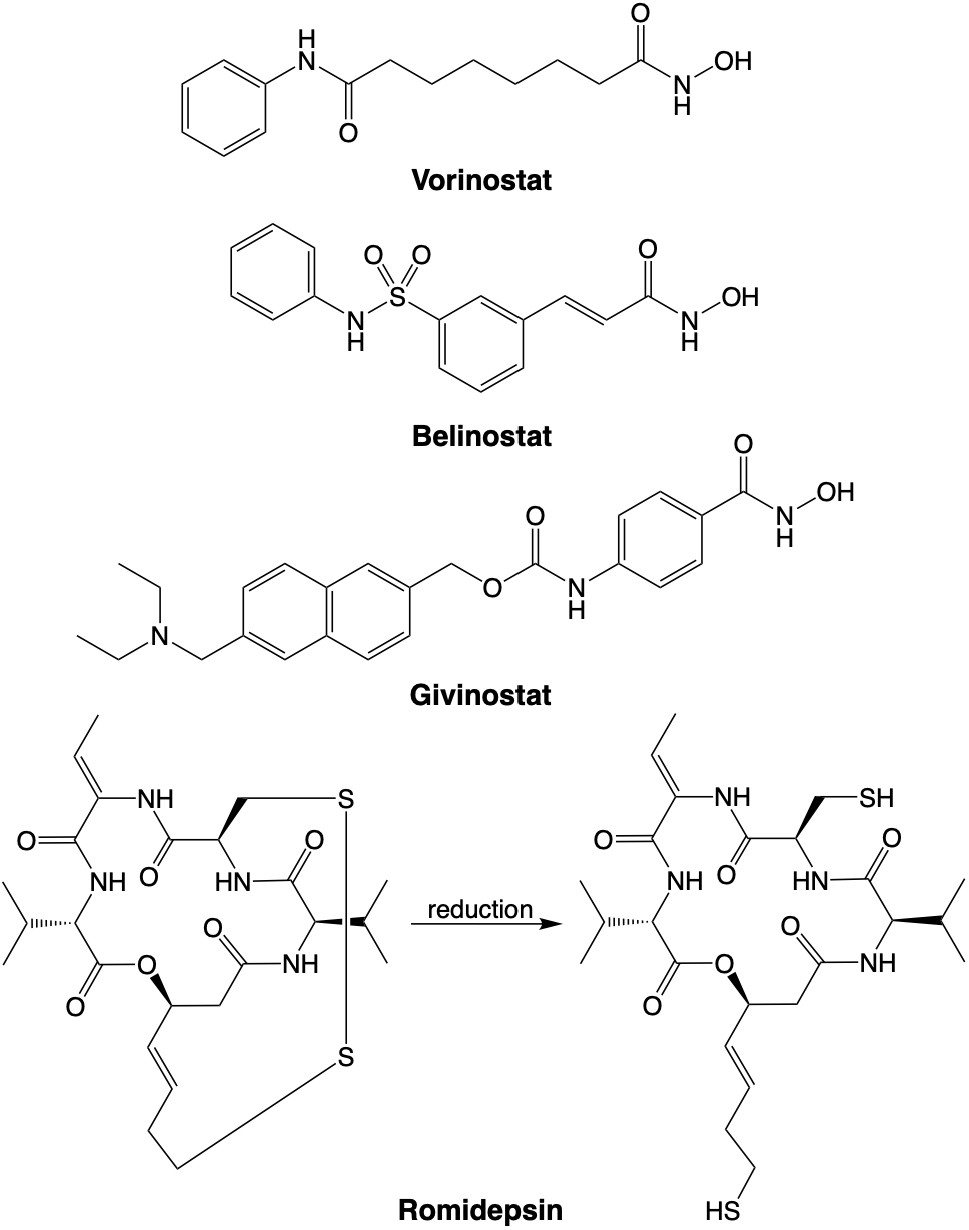
Molecular structures of HDAC inhibitors approved for clinical use by the U.S. Food and Drug Administration. Romidepsin undergoes disulfide reduction *in vivo* to yield the active form of the inhibitor with a thiol zinc-binding group.

Cyclic peptide natural products represent an exciting class of compounds in drug development and occupy a vast expanse of chemical space.^42,43^ Indeed, the cyclic tetrapeptide inhibitor trapoxin A (Figure 2)^44^ was instrumental in the development of an affinity column that enabled the isolation and discovery of human HDACs.^45^ Cyclic tetrapeptide inhibitors related to trapoxins A and B include HC toxin^46^ and apicidin.^47^ All of these are mechanism-based HDAC inhibitors: each contains a residue bearing a reactive ketone side chain that undergoes nucleophilic attack by zinc-bound water in the active site.^29,38,48,49^ Cyclic peptide natural products also include cyclic depsipeptides, macrocycles that contain both ester and amide linkages, such as the FDA-approved drug Romidepsin^50,51^ (Figure 1) and the structurally-related marine natural product Largazole (Figure 2).^52^ While the crystal structure of an HDAC-Romidepsin complex is not available, the crystal structure of the HDAC8-Largazole complex reveals that a potent thiolate-zinc coordination interaction dominates inhibitor binding.^53^

**Figure 2.**
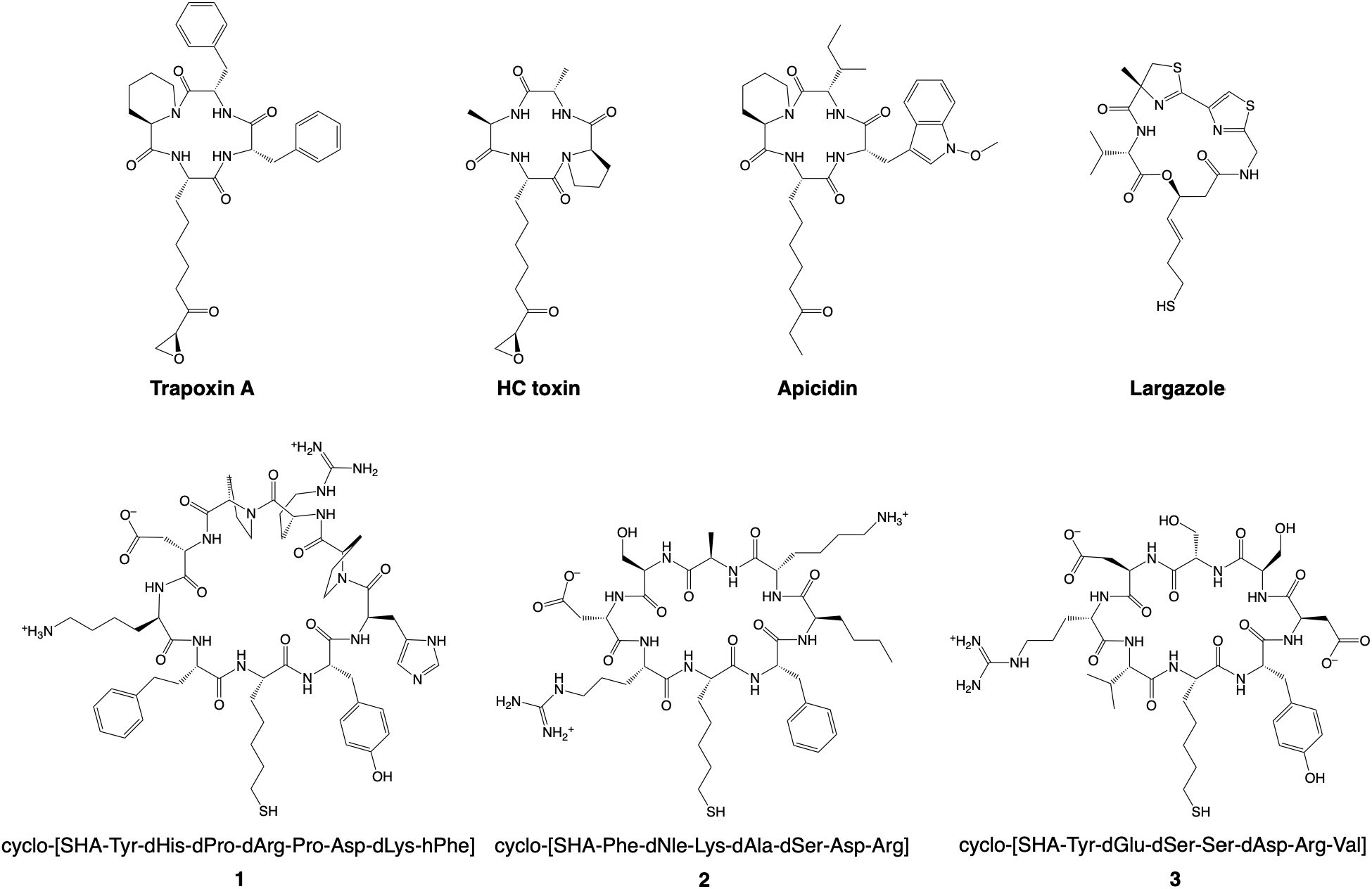
(Top) Cyclic peptide natural product inhibitors of HDACs. The marine natural product Largazole is a cyclic depsipeptide that undergoes thioester hydrolysis to yield the active form of the inhibitor as shown. (Bottom) Macrocyclic nonapeptide **1** inhibits HDAC6 with IC_50_ = 22 nM. Macrocyclic octapeptide **2** inhibits HDAC6 with IC_50_ = 17 nM. Macrocyclic octapeptide **3** inhibits HDAC6 with IC_50_ = 4.4 nM. Abbreviations: SHA = (S)-2-amino-7-sulfanylheptanoic acid, hPhe = homophenylalanine, Nle = norleucine; “d” indicates a D-amino acid.^56^ In the current work, residues in cyclic peptides are designated by three-letter abbreviations and sequence number, beginning with SHA-1; enzyme residues are designated by one-letter abbreviations and sequence number.

Cyclic peptide natural products readily mimic structural features of proteins, potentially facilitating ideal complementarity with the surface of a target protein.^54,55^ The same is true for *de novo* designed cyclic peptides containing both canonical and noncanonical amino acids. For example, computationally designed cyclic octapeptides and nonapeptides bind to HDACs with nanomolar inhibitory potencies and moderate isozyme selectivities.^56^ The design approach, termed “anchor extension”, involves docking the noncanonical amino acid (*S*)-2-amino-7-sulfanylheptanoic acid (SHA) in an HDAC active site followed by extending and cyclizing the bound peptide *in silico*. Crystal structures of HDAC6 complexed with cyclic octapeptide inhibitors generated in this study validated the success of this approach.^56,57^

Here, we report the 1.9 Å-resolution crystal structure of cyclic nonapeptide **1** (IC_50_ = 22 nM; Figure 2)^56^ complexed with HDAC6 CD2, showing that a potent thiolate-zinc coordination interaction anchors the cyclic peptide in the active site. Additionally, direct and water-mediated hydrogen bonds orient and stabilize the bound macrocycle. Surprisingly, the crystalline HDAC6 CD2–**1** complex was prepared by soaking pre-formed HDAC6 crystals in buffer solutions containing the 1,215-Dalton peptide, which thus is not too large to diffuse through solvent channels in the crystal. Thus, crystal soaking is a viable approach for the preparation of HDAC6-inhibitor complexes that are refractory to cocrystallization, even for exceptionally large inhibitors.

## Materials and Methods

### General

All chemicals and reagents were purchased from Sigma Aldrich and used without further purification. The synthesis and characterization of **1**, as well as its inhibitory potencies against HDAC6 and HDAC2, were previously reported.^56^

### Expression, purification, and crystallization of HDAC6 CD2

Catalytic domain 2 from *Danio rerio* (zebrafish) HDAC6 (UniProt F8W4B7) was encoded in the His6-MBP-TEV-HDAC6-pET28a(+) vector and expressed in *Escherichia coli* BL21(DE3) cells and purified as described previously.^58^ For crystallization, protein solution [10 mg/mL HDAC6 CD2 in 50 mM 4-(2-hydroxyethyl)-1-piperazineethanesulfonic acid (HEPES) (pH 7.5), 100 mM KCl, 5% glycerol, 1 mM tris(2-carboxyethyl)phosphine (TCEP)] was filtered with a 0.22-µm centrifuge filter. To crystallize unliganded HDAC6 CD2 using the sitting-drop vapor diffusion method at 13°C, 2 µL of protein solution were added to 1 µL of precipitant solution [0.1 M bis-tris (pH 6.0), 0.2 M KSCN, 20% polyethylene glycol (PEG) 3350] and equilibrated against a 500-µL reservoir of precipitant solution. Crystals typically appeared within 3 days and were transferred to a fresh sitting-drop vapor diffusion plate in wells containing 3 µL of inhibitor solution [0.1 M bis-tris (pH 6.0), 0.2 M KSCN, 20% polyethylene glycol 3350, 0.5 mM **1**] and equilibrated against a 500-µL reservoir of precipitant solution. After soaking for 2.5 days at 13°C, the inhibitor solution was replaced with a cryoprotectant solution [0.1 M bis-tris (pH 6.0), 0.2 M KSCN, 20% PEG 3350, 20% ethylene glycol (EGOH)]. Crystals were harvested and flash-cooled in liquid nitrogen.

### Data collection, structure determination, and refinement

X-ray diffraction data for the crystalline HDAC6 CD2-**1** complexes were collected remotely at the National Synchrotron Light Source II (NSLS-II), beamline 17-ID-2 (Brookhaven National Laboratory, Upton, NY). All diffraction data were indexed, integrated, scaled, and merged using XDS (ver. BUILT 20241002),^59^ POINTLESS (ver. 1.12.14),^60^ and AIMLESS (ver. 0.7.9)^61^ as part of the autoPROC (ver. 1.0.5)^62^ pipeline, and subjected to anisotropy analysis with STARANISO (ver. 2.4.9).^63^ Data were moderately anisotropic, with diffraction maxima observed at 1.8–1.9 Å resolution along the *a*- and *c*-axes, but only 2.48 Å along the *b*-axis. To generate an initial electron density map of the HDAC6 CD2-**1** complex, the structure of cyclohexenylhydroxamate-bound HDAC6 (PDB 6CSP) was used as a molecular replacement probe with Phaser-MR.^64^ The data yielded satisfactory molecular replacement solutions with two chains in the asymmetric unit. Iterative model building and refinement were performed using Coot (ver. 0.9.8.7 EL)^65^ and Phenix (ver. 1.21.2-5419-000),^66,67^ respectively. Inhibitor **1** binds only in the active site of chain B; a Zn^2+^-bound acetate ion is observed in the active site of chain A. The final structure was validated with MolProbity.^68^ Data collection and refinement statistics are recorded in Table 1.

**Table 1.** Data Collection and Refinement Statistics for the HDAC6–1 Complex^a^.

| **Space group** | *P*1 |
|---|---|
| ***a,b,c* (Å)** | 48.17, 55.24, 76.00 |
| **α, β, γ (°)** | 68.89, 87.38, 82.66 |
| **R<sub>merge</sub><sup>b</sup>** | 0.150 (0.518) |
| **CC<sub>1/2</sub><sup>c</sup>** | 0.954 (0.322) |
| **Redundancy** | 1.7 (1.5) |
| **Completeness (%)** | 87.8 (60.2) |
| **I/σ** | 2.0 (0.86) |
| **Refinement** |  |
| **Resolution (Å)** | 32.98 – 1.90<br>(1.95 – 1.90) |
| **No. reflections** | 50363 (2481) |
| **R<sub>work</sub>/R<sub>free</sub><sup>d</sup>** | 0.193 (0.291)/0.234 (0.342) |
| **Number of Atoms** |  |
| **Protein (2 chains)** | 5360 |
| **Ligand** | 155 |
| **Solvent** | 228 |
| **Average B factors (Å<sup>2</sup>)** |  |
| **Protein** | 14 |
| **Ligand** | 36 |
| **Solvent** | 16 |
| **Root-mean-square deviations** |  |
| **Bond lengths (Å)** | 0.002 |
| **Bond angles (°)** | 0.6 |
| **Ramachandran plot<sup>e</sup>** |  |
| **Favored (%)** | 96.83 |
| **Allowed (%)** | 3.03 |
| **Outliers (%)** | 0.14 |
| **MolProbity score** | 0.29 |
| **PDB code** | 39GH |

<sup>a</sup>Values in parentheses refer to the highest-resolution shell of data. <sup>b</sup>R<sub>merge</sub> = ∑<sub>h</sub>∑<sub>i</sub>|I<sub>i,h</sub> − ⟨I⟩<sub>h</sub>|/∑<sub>h</sub>∑<sub>i</sub>I<sub>i,h</sub>, where ⟨I⟩<sub>h</sub> is the average intensity calculated for reflection h from i replicate measurements. <sup>c</sup>Pearson correlation coefficient between random half-datasets. <sup>d</sup>R<sub>work</sub> = ∑||F<sub>o</sub>| − |F<sub>c</sub>||/∑|F<sub>o</sub>| for reflections contained in the working set. |F<sub>o</sub>| and |F<sub>c</sub>| are the observed and calculated structure factor amplitudes, respectively. R<sub>free</sub> is calculated using the same expression for reflections contained in the test set held aside during refinement. <sup>e</sup>Calculated with MolProbity.

## Results

The X-ray crystal structure of the HDAC6 CD2–**1** complex was solved at 1.9-Å resolution (Figure 3). Electron density for the macrocyclic nonapeptide is continuous and well-defined for zinc-bound SHA-1 (S•••Zn^2+^ distance = 2.3 Å) and neighboring amino acids in the macrocycle, but there are some breaks in density, as well as absent density for much of the Pro-6–Asp-7 segment, in the portion of the macrocycle that extends into a large solvent-filled region of the crystal lattice. Broken or missing electron density in this region is likely due to conformational disorder of residues that are not stabilized by binding to the enzyme. All peptide linkages in well-defined electron density adopt *trans* conformations.

**Figure 3.**
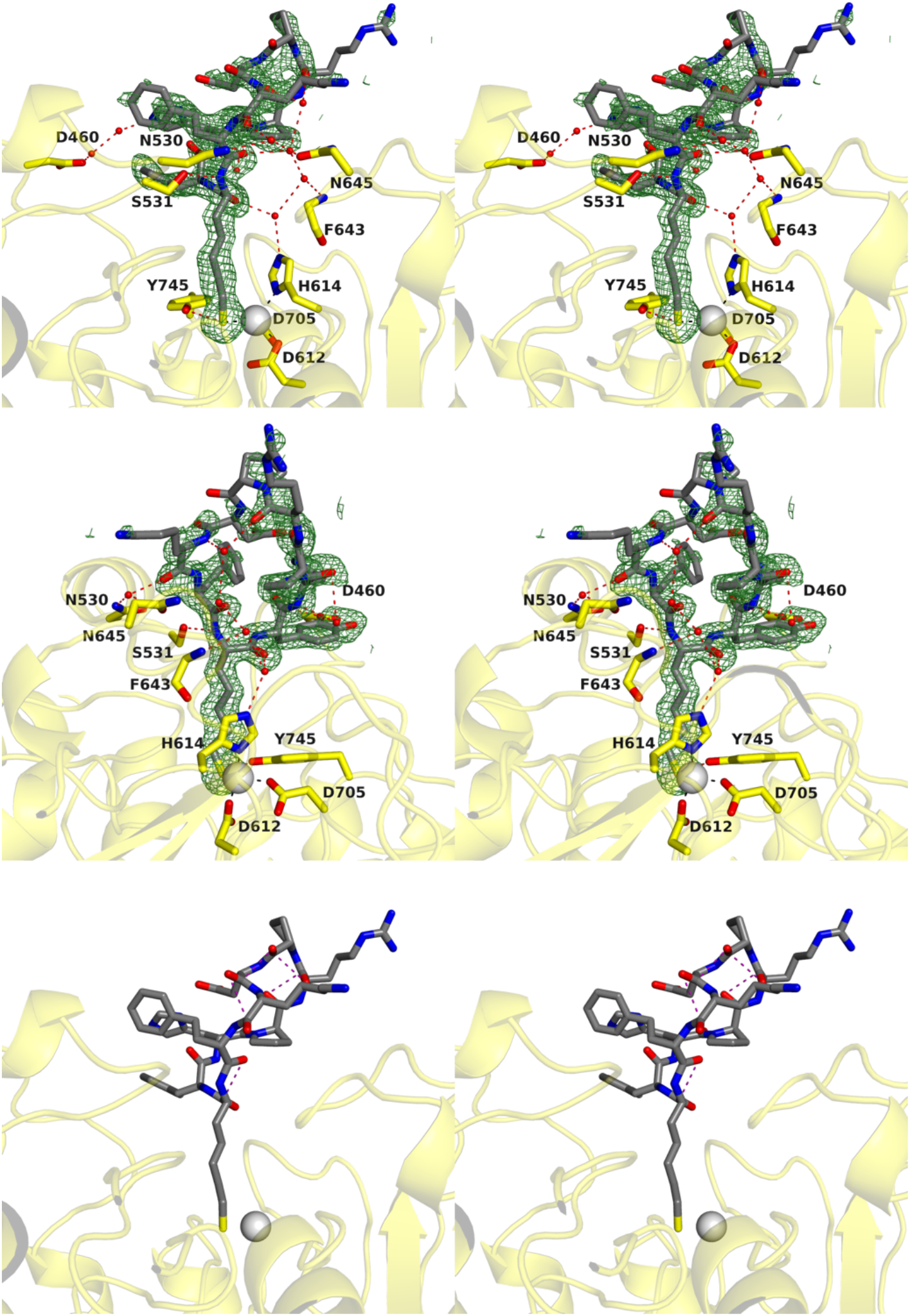
(Top and middle) Stereoviews of two perpendicular orientations of a Polder omit map (contoured at 2.7 σ) showing the binding of cyclic nonapeptide **1**, cyclo-SHA-Tyr-dHis-dPro-dArg-Pro-Asp-dLys-hPhe (SHA = (S)-2-amino-7-sulfanylheptanoic acid, hPhe = homophenylalanine; “d” designates a D-amino acid), in the active site of HDAC6 CD2. (Bottom) Stereoview of cyclic nonapeptide inhibitor 1 bound to HDAC6 with n→π* interactions highlighted. Atomic color coding: C = gray (**1**), yellow (HDAC6); N = blue; O = red; S = dark yellow; Zn^2+^ = gray. Interaction color coding: black (metal coordination); red (hydrogen bonds); purple (n→π* interactions). Other: red spheres (water), green mesh (polder map).

Cyclic nonapeptide **1** makes only two direct hydrogen bonds in the enzyme active site: the zinc-bound thiolate sulfur of SHA-1 accepts a hydrogen bond from the phenolic hydroxyl group of the catalytic tyrosine, Y745; the backbone NH group of SHA-1 donates a hydrogen bond to the hydroxyl group of S531. Additionally, water-mediated enzyme-inhibitor hydrogen bonds are observed: the backbone carbonyl and NH groups of SHA-1, dArg-5, dLys-8, and hPhe-9 accept hydrogen bonds from water molecules that, in turn, form hydrogen bonds with the imidazole group of Zn^2+^ ligand H614 as well as the backbone NH group of F643. Although the density is relatively weak, there is a pair of water molecules that for a set of hydrogen atoms bridging nonapeptide residue dHis-3 and HDAC6 residue D460 on the opposite side of the active site entrance. No intramolecular hydrogen bonds involving residues with well-defined electron density are observed within the nonapeptide macrocycle. Hydrogen bonds are observed between the side chains of Tyr-2 and T777, and between the backbone carbonyls of dHis-3 and dPro-4 and the side chain of N784 in a separate copy of HDAC6 CD2 in a neighboring unit cell.

Five intramolecular *n*→π* interactions are observed in cyclic nonapeptide **1** (Figure 3). These interactions occur between the carbonyl oxygen of residue *n* and the carbonyl carbon of residue *n* + 1 in the following residue pairs, respectively: hPhe-9 and SHA-1 (O•••C separation = 3.2 Å), dPro-4 and dArg-5 (O•••C separation = 3.0 Å), dArg-5 and Pro-6 (O•••C separation = 3.1 Å), Pro-6 and Asp-7 (O•••C separation = 2.7 Å), and Asp-7 and dLys-8 (O•••C separation = 3.3 Å). Interestingly, intramolecular *n*→π* interactions occur between residues *n* and *n*+1 in α-helices;^69,70^ indeed, the Φ and Ψ angles of SHA-1 correspond to those of a residue in an α-helix. There are no intermolecular *n*→π* interactions observed between cyclic nonapeptide **1** and HDAC6.

All residues in the cyclic nonapeptide adopt allowed backbone torsion angles on a Ramachandran plot except for dHis-3, dPro-4, dArg5, and dLys8, which are all D-amino acids (Figure 4, Table 2). The Ramachandran plot was originally formulated for polypeptides containing L-amino acids,^71^ so variations are not unexpected for peptides containing mixtures of L- and D-amino acids. The backbone torsion angles of **1** most closely align with backbone torsion angles of SHA-1 in cyclic octapeptides **2** and **3** (Figure 2) bound in the HDAC6 CD2 active site, which most closely align with backbone torsion angles for acetyllysine substrates bound in the HDAC6 CD2 active site.^56,57^ The conformations of adjacent residues and the remainder of the cyclic peptide tend to diverge, some substantially so (Figure 4, Table 2).

**Figure 4.**
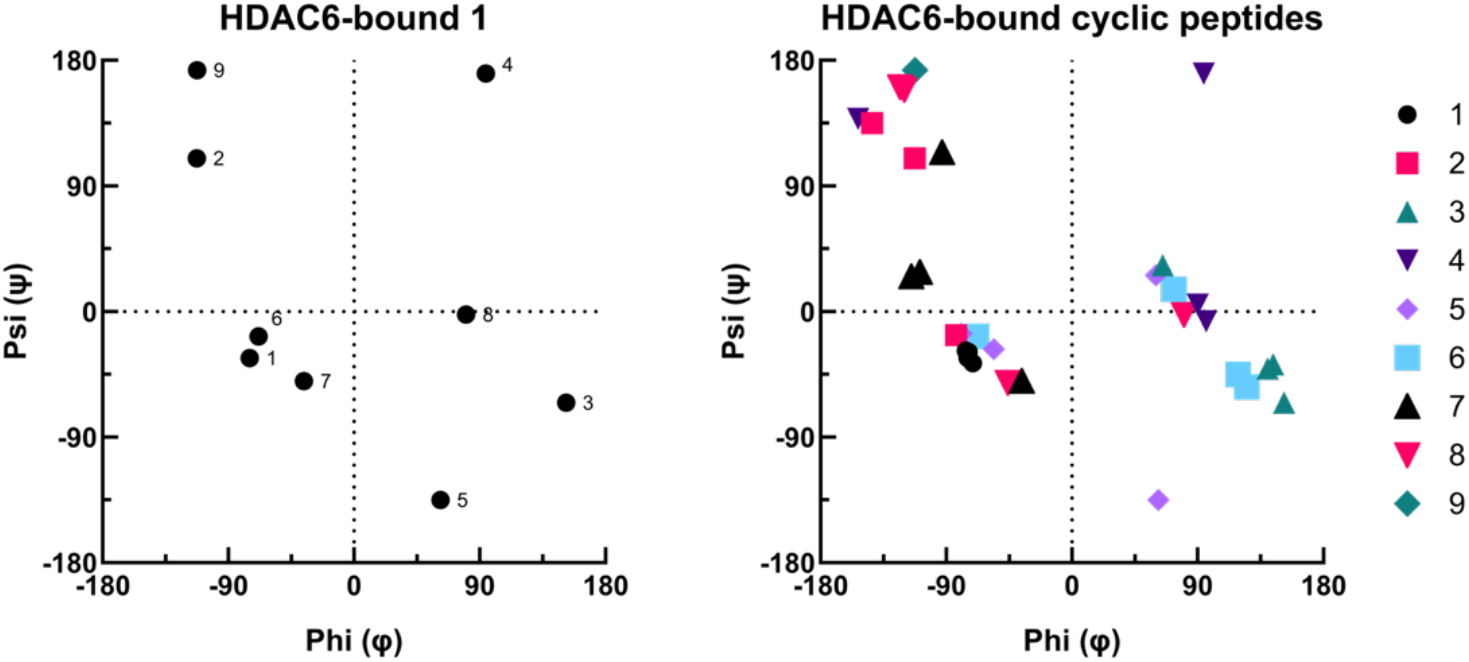
(Left) Ramachandran plot for cyclic nonapeptide **1** in the HDAC6 CD2-**1** complex. Residue numbering is defined in Table 2. (Right) Ramachandran plot for cyclic nonapeptide **1** and cyclic octapeptides **2** and **3** in their HDAC6 CD2 complexes. The Zn^2+^-bound SHA residue of each peptide is designated as residue 1.

**Table 2.** Backbone torsion angles (°) for cyclic peptide inhibitors bound to HDAC6 CD2.

| **Compound<sup>a</sup>** | **PDB<sup>b</sup>** |  | **1** | **2** | **3** | **4** | **5** | **6** | **7** | **8** | **9** |
|---|---|---|---|---|---|---|---|---|---|---|---|
| **(1)** | **39GH** | residue | SHA | Tyr | dHis | dPro | dArg | Pro | Asp | dLys | hPhe |
|  |  | phi (ϕ) | -75 | -113 | 152 | 94 | 62 | -68 | -36 | 80 | -112 |
|  |  | psi (ψ) | -33 | 110 | -65 | 170 | -135 | -18 | -50 | -2 | 173 |
| **(2)** | **6WSJ** | residue | SHA | Phe | dNle | Lys | dAla | dSer | Asp | Arg | — |
|  |  | phi (ϕ) | -71 | -83 | 65 | -153 | 60 | 73 | -93 | -46 |  |
|  |  | psi (ψ) | -37 | -17 | 33 | 138 | 26 | 16 | 114 | -51 |  |
| **(3) (monomer A)** | **8EQI** | residue | SHA | Tyr | dGlu | dSer | Ser | dAsp | Arg | Val | — |
|  |  | phi (ϕ) | -74 | -143 | 140 | 96 | -56 | 119 | -115 | -120 |  |
|  |  | psi (ψ) | -29 | 135 | -41 | -7 | -27 | -45 | 25 | 159 |  |
| **(3) (monomer B)** | **8EQI** | residue | SHA | Tyr | dGlu | dSer | Ser | dAsp | Arg | Val | — |
|  |  | phi (ϕ) | -76 | -143 | 144 | 90 | -79 | 125 | -109 | -123 |  |
|  |  | psi (ψ) | -28 | 135 | -38 | 5 | -16 | -54 | 28 | 161 |  |

<sup>a</sup>Cyclic peptides **1**, **2**, and **3** initially designated des3.2.2, des4.3.1, and des4.2.0, respectively, in ref. 55.
<sup>b</sup>Crystal structures of HDAC6 CD2 complexes with compounds **2** and **3** reported in refs. 55 and 56, respectively.

## Discussion

Because we were unsuccessful in cocrystallizing HDAC6 CD2 with cyclic nonapeptide **1**, the preparation of the crystalline HDAC6 CD2–**1** complex required soaking pre-formed crystals of unliganded HDAC6 CD2 in a buffer solution containing 0.5 mM **1** for 2.5 days. The crystal soaking approach for preparing protein complexes with small organic ligands was first employed in the preparation of lysozyme complexes with N-acetylglucosamine derivatives.^72^ Since then, this approach has been a valuable complement to cocrystallization, and especially so for complexes that do not readily cocrystallize. On average, protein crystals contain ∼50% solvent, so ligands can diffuse through solvent channels to bind to protein functional sites as long as these binding sites are not sterically occluded. Remarkably, rates of ligand diffusion into a crystalline protein vary over several orders of magnitude and can take from tenths of a second to a few months to achieve binding with occupancy sufficient for detection by protein crystallography.^73^

However, crystal soaking is not necessarily superior to cocrystallization if the target protein undergoes significant conformational changes upon ligand binding that might be hindered in the crystal lattice.^74^ The success of the crystal soaking approach further depends on the size and steric bulk of the ligand, which of course must be sufficiently small and flexible to diffuse through constricted solvent channels in the crystal. Accordingly, crystal soaking experiments are most often successful with small molecule ligands no more than a few hundred Daltons in size. A notable exception is the preparation of crystalline tropomyosin complexed with a 19-kDa fragment of troponin soaked into crystals – the large unit cell of tropomyosin and 95% solvent content facilitated structure determination of the tropomyosin-troponin fragment complex, albeit at 17 Å resolution.^75^

Numerous crystal structures of HDAC6 CD2-inhibitor complexes prepared by cocrystallization reveal minimal protein structural changes triggered by inhibitor binding.^29,56,57^ The HDAC6 CD2 crystals used here for soaking experiments contain 47% solvent; the active site of chain B in the *P*1 unit cell opens up to a large, solvent-filled cavity in the crystal lattice and is sufficiently accessible for inhibitor binding. Thus, HDAC6 CD2 is suitable for crystal soaking experiments, and we have already demonstrated the success of this approach for much smaller inhibitors with molecular weights ranging from 412 to 662 Da.^76^ Here, we demonstrate the robustness of the crystal soaking approach with a significantly larger inhibitor that yields a 1.9-Å resolution crystal structure of the enzyme-inhibitor complex.

The X-ray crystal structure of the HDAC6 CD2-**1** complex confirms the general binding mode predicted from the *in silico* anchor extension approach.^56^ Interestingly, there are only three direct enzyme-inhibitor hydrogen bonds made with **1**, two of which are made with the Zn^2+^-bound SHA-1 residue. The remainder of the cyclic nonapeptide provides only one direct and two water-mediated enzyme-inhibitor hydrogen bonds, yet these interactions must contribute to 30-fold enhanced inhibitory potency of **1** (IC_50_ value = 22 nM) compared to pure SHA (IC_50_ = 670 nM).^56^ Comparable enhancement of inhibitory activity is observed for cyclic octapeptide **2** (IC_50_ = 17 nM), and even greater enhancement (∼150-fold) is observed for cyclic octapeptide **3** (IC_50_ = 4.4 nM). The structural basis for the five-fold difference in inhibitory potencies of **1** and **3** is not clear.

Structural comparisons of the HDAC6 CD2 complexes with cyclic peptides **1**–**3** yield useful insight. Each inhibitor is anchored in the active site by its SHA-1 residue, which makes a potent thiolate-Zn^2+^ coordination interaction. The backbone torsion angles of SHA-1 are quite similar in each complex (Figure 4); however, residue conformations tend to diverge more the further away they are from SHA-1 (Figure 5a). Interestingly, an overlay of cyclic tetrapeptide, octapeptide, and nonapeptide inhibitors illustrates the increasing span of the inhibitors as the number of amino acids in the macrocycle increases (Figure 5b). At the most distant point on the main chain backbone, HC toxin tetrapeptide, octapeptide **3**, and nonapeptide **1** extend 15 Å, 17 Å, and 18 Å, respectively, away from the Zn^2+^ ion at the bottom of the active site. Similarly, nonapeptide **1** extends across a large portion of the surface of the active site. At the furthest points of the side chains on Tyr-2 and dLys-8, the inhibitor spans 20 Å, allowing it to make indirect hydrogen bonds with both residues D640 and N645, despite their significant spatial separation (Figure 3).

**Figure 5.**
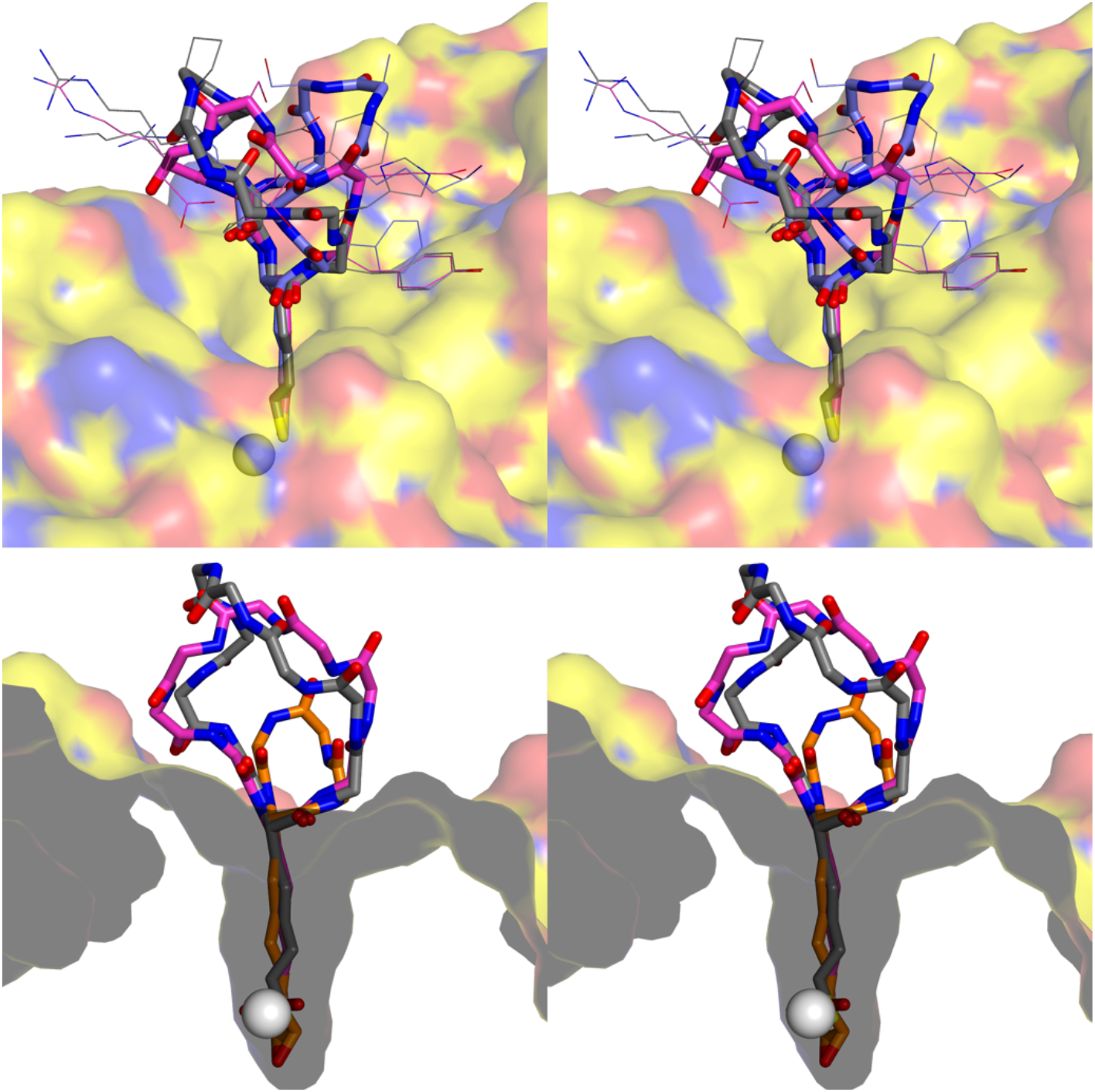
(Top) Stereoscopic overlay of **1**, **2** (PDB 6WSJ), and **3** (PDB 8EQI) bound in the active site of HDAC6 CD2. (Bottom) Stereoview, rotated ∼90° from (a), of the backbone binding conformations of nonapeptide **1**, compared with that of octapeptide **3** and the tetrapeptide HC toxin (PDB 5EFJ). Molecular structures of all cyclic peptides are shown in Figure 2. Atomic color coding: C = gray (**1**), slate (**2**), magenta (**3**), orange (HC toxin), yellow (protein surface); N = blue; O = red; S = dark yellow.

For each of these cyclic peptide inhibitors, there are differences in intramolecular and intermolecular interactions, but intramolecular and intermolecular interactions of zinc-bound SHA-1 are conserved across all three structures: the zinc-bound thiolate of SHA-1 accepts a hydrogen bond from catalytic tyrosine Y745, the backbone NH group of SHA-1 donates a hydrogen bond to S531; and the backbone carbonyl group of the preceding residue (**1**: hPhe-9; **2**: Arg-8; **3**: Val-8) engages in an *n*→π* interaction with the backbone carbonyl group of SHA-1. As previously mentioned, *n*→π* interactions between immediately adjacent residues comprise a stabilizing feature of α-helices.^69,70^

Water molecules near the active site of HDAC6 CD2 form a hydrogen bonded network involving **1**, the backbone NH group of F643, and zinc ligand H614. In the crystal structure of the HDAC6 CD2-**2** complex, the hydroxyl groups of a single molecule of ethylene glycol occupy the same space as two of the water molecules seen in the structure of the HDAC6 CD2-**1** complex. Thus, the cryoprotectant molecule can substitute for water molecules in forming bridging hydrogen bond interactions between the enzyme and the bound inhibitor (Figure 6).

**Figure 6.**
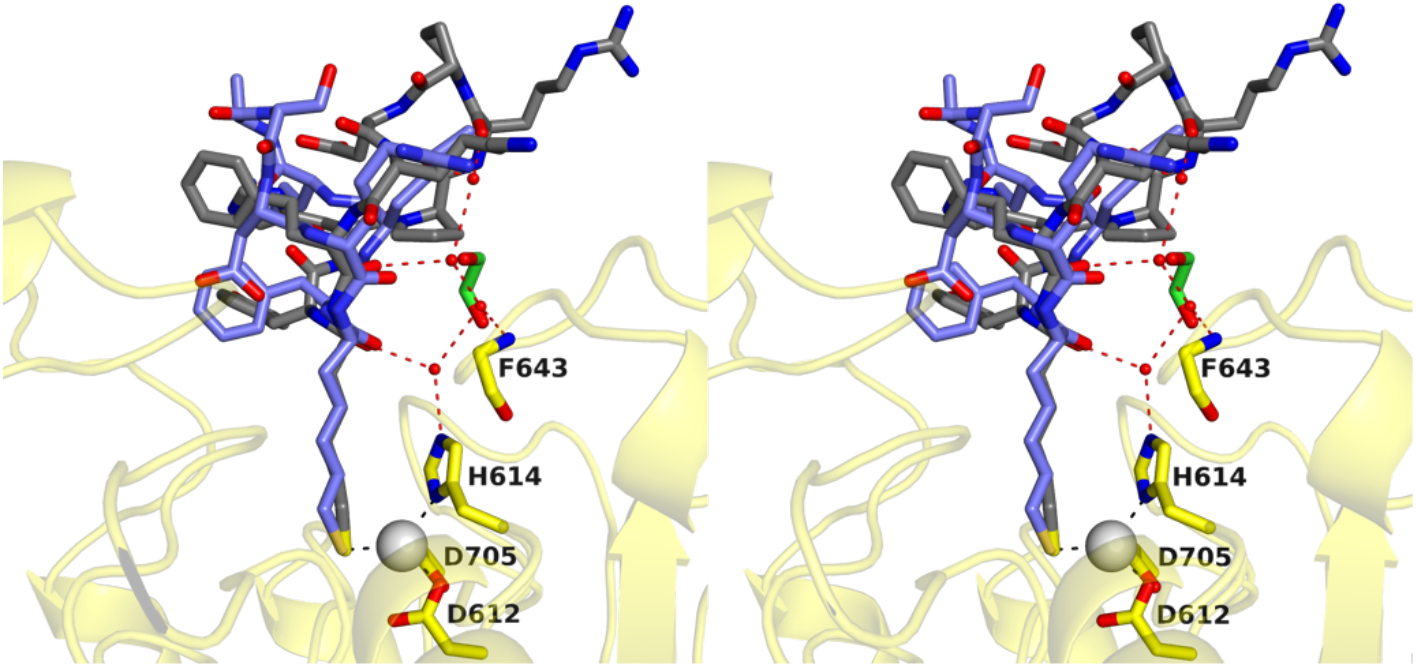
Stereoscopic overlay of **1** and **2** bound in the active site of HDAC6 CD2. Atomic color coding: C = gray (**1**), slate (**2**), green (ethylene glycol), yellow (protein); N = blue; O = red; S = dark yellow.

All three inhibitors make direct interactions with S531 through the backbone NH group of SHA-1. All three inhibitors also make indirect interactions with N530 through a hydrogen bonded water molecule. This water molecule bridges the gap between the carboxamide carbonyl group of N530 and the backbone NH group of dLys-8 in **1**, Arg-8 in **2**, and the carbonyl group of Arg-7 in **3**. This same water molecule also bridges the gap between N530 and N645, further stabilizing these residues in place. N530, which corresponds to D567 in the human enzyme, is a key residue in the active site of HDAC6. This residue is located at the mouth of the active site and is one of the few active site residues not conserved in other HDAC isozymes. Targeting D567 in human HDAC6 CD2 could be an avenue toward developing isozyme-selective inhibitors with suitably positioned hydrogen bond donors capable of interacting with D567. For example, BAS-2, an HDAC6 CD2-selective mechanism-based inhibitor,^77^ is active against wild-type human HDAC6 and N530D zebrafish HDAC6, but not D567N human HDAC6 or wild-type zebrafish HDAC6.^78^ In this example, D567 of human HDAC6 serves as a gatekeeper for inhibitor binding and activation. In view of the growing prominence of D567 for inhibitor binding and activation in human HDAC6, a “humanized” version of zebrafish HDAC6 CD2 containing the N530D substitution should be used to evaluate the importance of enzyme-inhibitor interactions in the outer active site cleft.

Finally, the binding mode of **1** in the active site of HDAC6 CD2 resembles that of an acetyllysine peptide substrate (Figure 7). The φ and ψ angles of SHA-1 in HDAC6 CD2-bound **1** are -75° and -33°, respectively. These torsion angles are comparable to those of a bound substrate, for which the φ and ψ angles are -73° and -38°, respectively (PDB 5EFK).^29^ Additionally, the structure of the HDAC6 CD2-bound acetyllysine peptide substrate reveals similar interactions with S531 via the acetyllysine residue in a similar fashion to SHA in cyclic nonapeptide **1**. This similarity demonstrates the ability of the inhibitor to effectively mimic a naturally occurring substrate.

**Figure 7.**
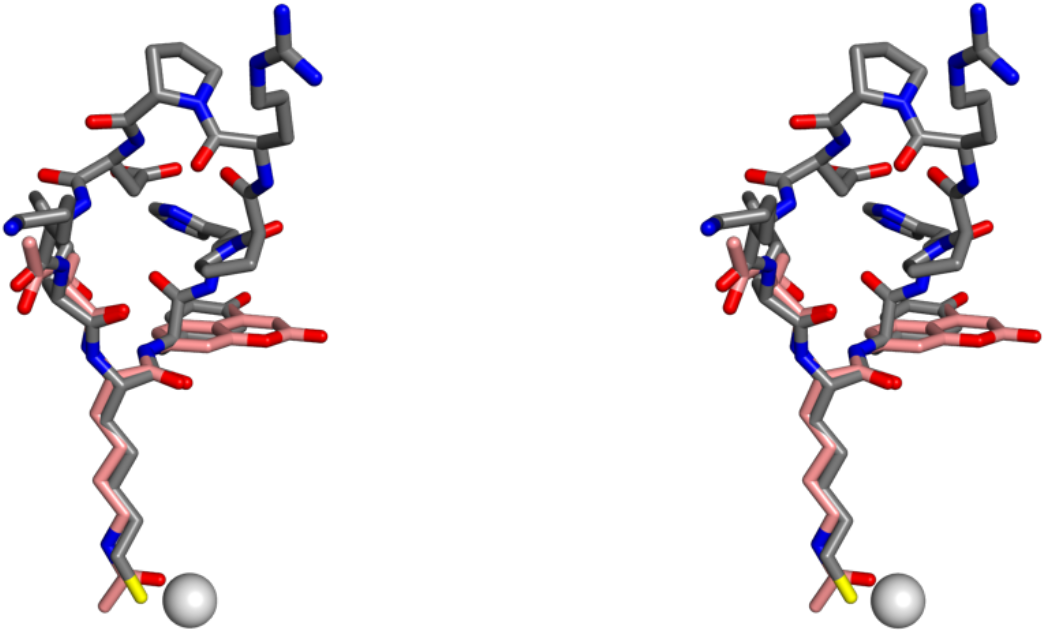
Stereoscopic overlay of **1** (gray) and α-tubulin K40 tripeptide substrate (salmon) bound in the active site of HDAC6 CD2. Atomic color coding: C = gray (**1**), salmon (substrate); N = blue; O = red; S = dark yellow. For reference, the Zn^2+^ ion is shown as a gray sphere.

## Concluding Remarks

On average, one cyclic peptide per year has been approved for clinical use during the past three decades.^79^ The complex three-dimensional shapes of cyclic peptides, their metabolic stabilities, their resistance to proteolytic degradation, and their wide range of applications continue to drive major drug discovery campaigns.^80^ The tricyclic macrocyclic peptide Enlicitide, a PCSK9 inhibitor recently approved by the U.S. Food and Drug Administration for the treatment of hypercholesterolemia, is a hallmark example.^81^ Both canonical and noncanonical amino acids can be incorporated into cyclic peptide designs, so these relatively small scaffolds can yield impressive chemodiversity in terms of their amino acid content and structure.

Although natural products still represent the predominant origin of cyclic peptide inhibitors in clinical use, there has been a recent surge in artificial intelligence- and computationally-driven strategies for *de novo* cyclic peptide design.^82^ These advances help to minimize the high cost, time consumption, and chemical space limitations of more traditional drug design strategies, though *in silico* hits must of course be validated experimentally.^83^ As macrocyclic peptide inhibitors continue to grow in ubiquity and computational capabilities continue to advance, the structure of the HDAC6 CD2-**1** complex with a macrocyclic nonapeptide inhibitor will serve as an important foundation for the optimization of a high-affinity cyclic peptide inhibitor.

## ASSOCIATED CONTENT

## Accession Code

Atomic coordinates and structure factor amplitudes for the HDAC6-**1** complex have been deposited in the Protein Data Bank (www.rcsb.org) with accession code 39GH.

## AUTHOR INFORMATION

## Authors

**Abraham Eafa –** Roy and Diana Vagelos Laboratories, Department of Chemistry, University of Pennsylvania, Philadelphia, Pennsylvania 19104-6323, United States.

**Parisa Hosseinzadeh –** Department of Bioengineering, Knight Campus, University of Oregon, Eugene, OR 97403 United States.

**David Baker –** Department of Biochemistry, Institute for Protein Design, University of Washington, Seattle, WA 98195 United States.

## ACKNOWLEDGMENTS

We thank Drs. Paris R. Watson, Juana Goulart Stollmaier, and Luigi F. Di Costanzo for helpful scientific discussions and guidance. This work is based on research conducted at beamline 17-ID-2 (FMX) of the National Synchrotron Light Source II, a DOE Office of Science User Facility operated for the DOE Office of Science by Brookhaven National Laboratory under Contract DE-SC0012704. The Center for BioMolecular Structure (CBMS) is primarily supported by the National Institutes of Health, NIGMS, through a Center Core P30 Grant (P30GM133893) and by the DOE Office of Biological and Environmental Research (KP1605010).

## References

1. Gershey, E. L., Vidali, G., Allfrey, V. G. (1968) Chemical studies of histone acetylation: the occurrence of ε-*N*-acetyllysine in the *f2a1* histone. J. Biol. Chem. 243, 5018–5022.

2. Vidali, G., Gershey, E. L., Allfrey, V. G. (1968) Chemical studies of histone acetylation: the distribution of ε-*N*-acetyllysine in calf thymus histones. J. Biol. Chem. 243, 6361–6366.

3. Lin, H., Su, X., He, B. (2012) Protein lysine acetylation and cysteine succination by intermediates of energy metabolism. ACS Chem. Biol. 7, 947–960.

4. Verdin, E., Ott, M. (2015) 50 years of protein acetylation: from gene regulation to epigenetics, metabolism and beyond. Nat. Rev. Mol. Cell Biol. 16, 258–264.

5. Zhang, Z., Tan, M., Xie, Z., et al. (2011) Identification of lysine succinylation as a new post-translational modification. Nat. Chem. Biol. 7, 58–63.

6. Alleyn, M., Breitzig, M., Lockey, R., Kolliputi, N. (2018) The dawn of succinylation: a posttranslational modification. Am. J. Physiol. Cell. Physiol. 314, C228–C232.

7. Giglione, C., Meinnel, T. (2022) Mapping the myristoylome through a complete understanding of protein myristoylation biochemistry. Prog. Lipid Res. 85, 101139.

8. Komaniecki, G., Lin, H. (2021) Lysine fatty acylation: regulatory enzymes, research tools, and biological function. Front. Cell Dev. Biol. 9, 717503.

9. Zhao, S., Zhang, X., Li, H. (2018) Beyond histone acetylation – writing and erasing histone acylations. Curr. Opin. Struct. Biol. 53, 169–177.

10. Shukri, A. H., Lukinović, V., Charih, F., Biggar, K. K. (2023) Unraveling the battle for lysine: a review of the competition among post-translational modifications. *Biochim. Biophys. Acta*, Gene Regul. Mech. 1866, 194990.

11. Hintzen, J. C. J., Burslem, G. M. (2026) Beyond acetylation: evaluating the emerging biology of lysine acylation. Cell Chem. Biol. 10.1016/j.chembiol.2026.07.009.

12. Delcuve, G., Khan, D., Davie, J. (2012) Roles of histone deacetylases in epigenetic regulation: emerging paradigms from studies with inhibitors. Clin. Epigenetics 4, 5.

13. Kouzarides, T. (2000) Acetylation: a regulatory modification to rival phosphorylation? EMBO J. 19, 1176–1179.

14. Choudhary, C., Weinert, B., Nishida, Y., et al. (2014) The growing landscape of lysine acetylation links metabolism and cell signalling. Nat. Rev. Mol. Cell Biol. 15, 536–550.

15. Tan, K., Rodríguez-Fernández, M. M., Keuler, T., et al. (2026) Crosstalk between NLRP3 signaling and histone deacetylases in inflammasome-driven diseases. J. Med. Chem. 69, 102–120.

16. Zhao, S. et al. (2010) Regulation of cellular metabolism by protein lysine acetylation. Science 327, 1000–1004.

17. Wang, Q. et al. (2010) Acetylation of metabolic enzymes coordinates carbon source utilization and metabolic flux. Science 327, 1004–1007.

18. Nakayasu, E. S., Burnet, M. C., Walukiewicz, H. E., et al. (2017) Ancient regulatory role of lysine acetylation in central metabolism. mBio 8, e01894–17.

19. Brownell, J. E., Allis, C. D. (1996) Special HATs for special occasions: linking histone acetylation to chromatin assembly and gene activation. Curr. Opin. Genet. Dev. 6, 176–184.

20. Marmorstein, R., Zhou, M.-M. (2014) Writers and readers of histone acetylation: structure, mechanism, and inhibition. Cold Spring Harb. Perspect. Biol. 6, a018762.

21. Shvedunova, M., Akhtar, A. (2022) Modulation of cellular processes by histone and non-histone protein acetylation. Nat. Rev. Mol. Cell Biol. 23, 329–349.

22. Seto, E., Yoshida, M. (2014) Erasers of histone acetylation: the histone deacetylase enzymes. Cold Spring Harb. Perspect. Biol. 6, a018713.

23. Christianson, D. W. (2026) Molecular structure and function of zinc-dependent histone deacetylases. Annu. Rev. Biochem. 95, 173–197.

24. Falkenberg, K., Johnstone, R. (2014) Histone deacetylases and their inhibitors in cancer, neurological diseases and immune disorders. Nat. Rev. Drug Discov. 13, 673–691.

25. Gregoretti, I. V., Lee, Y. M., Goodson, H. V. (2004) Molecular evolution of the histone deacetylase family: functional implications of phylogenetic analysis. J. Mol. Biol. 338, 17–31.

26. Yuan, H., Marmorstein, R. (2012) Structural basis for sirtuin activity and inhibition. J. Biol. Chem. 287, 42428–42435.

27. Grozinger, C. M., Hassig, C. A., Schreiber, S. L. (1999) Three proteins define a class of human histone deacetylases related to yeast Hda1p. Proc. Natl. Acad. Sci. USA 96, 4868–4873.

28. Verdel, A., Khochbin, S. (1999) Identification of a new family of higher eukaryotic histone deacetylases. J. Biol. Chem. 274, 2440–2445.

29. Hai, Y., Christianson, D. (2016) Histone deacetylase 6 structure and molecular basis of catalysis and inhibition. Nat. Chem. Biol. 12, 741–747.

30. Kutil, Z., Skultetyova, L., Rauh, D., et al. (2019) The unraveling of substrate specificity of histone deacetylase 6 domains using acetylome peptide microarrays and peptide libraries. FASEB J. 33, 4035–4045.

31. Haggarty, S. J., Koeller, K. M., Wong, J. C., et al. (2003) Domain-selective small-molecule inhibitor of histone deacetylase 6 (HDAC6)-mediated tubulin deacetylation. Proc. Natl. Acad. Sci. USA 100, 4389–4394.

32. Cohen, T. J., Guo, J. L., Hurtado, D. E., et al. (2011) The acetylation of tau inhibits its function and promotes pathological tau aggregation. Nat. Commun. 2, 252.

33. Zhang, X., Yuan, Z., Zhang, Y., et al. (2007) HDAC6 modulates cell motility by altering the acetylation level of cortactin. Mol. Cell 27, 197–213.

34. Jung, M., Hoffmann, K., Brosch, G., Loidl, P. (1997) Analogues of trichostatin A and trapoxin B as histone deacetylase inhibitors. Bioorg. Med. Chem. Lett. 7, 1655–1658.

35. Ma, N., Luo, Y., Wang, Y., et al. (2016) Selective histone deacetylase inhibitors with anticancer activity. Curr. Top. Med. Chem. 16, 415–426.

36. Vickers, C. J., Olsen, C. A., Leman, L. J., Ghadiri, M. R. (2012) Discovery of HDAC inhibitors that lack an active site Zn^2+^-binding functional group. ACS Med. Chem. Lett. 3, 505–508.

37. Marcaurelle, L. A., Comer, E., Dandapani, S., et al. (2010) An aldol-based build/couple/pair strategy for the synthesis of medium- and large-size rings: discovery of macrocyclic histone deacetylase inhibitors. J. Am. Chem. Soc. 132, 16962–16976.

38. Beshore, D. C., Adam, G. C., Barnard, R. J. O., et al. (2021) Redefining the histone deacetylase inhibitor pharmacophore: high potency with no zinc cofactor interaction. ACS Med. Chem. Lett. 12, 540–547.

39. Lechner, S., Malgapo M. I. P., Grätz, C., et al. (2022) Target deconvolution of HDAC pharmacopoeia reveals MBLAC2 as common off-target. Nat. Chem. Biol. 18, 812–820.

40. Li, T., Zhang, C., Hassan, S., et al. (2018) Histone deacetylase 6 in cancer. J. Hematol. Oncol. 11, 111.

41. Shen, S., Kozikowski, A. (2016) Why hydroxamates may not be the best histone deacetylase inhibitors—what some may have forgotten or would rather forget? ChemMedChem 11, 15–21.

42. Bergamaschi, G., Lodigiani, G., Gandolfi, S., et al. (2026) Nature-inspired macrocyclic peptides: discovery and molecular engineering for drug development. Curr. Opin. Chem. Biol. 94, 102738.

43. Thite, T., Iyer, K. A., Jain, P., et al. (2026) Cyclic peptides in modern drug discovery: trends and therapeutic directions. J. Med. Chem. 69, 17770–17799.

44. Itazaki, H., Nagashima, K., Sugita, K., et al. (1990) Isolation and structural elucidation of new cyclotetrapeptides, trapoxins A and B, having detransformation activities as antitumor agents. J. Antibiot. 43, 1524–1532.

45. Taunton, J., Hassig, C. A., and Schreiber, S. L. (1996) A mammalian histone deacetylase related to the yeast transcriptional regulator Rpd3p. Science 272, 408–411.

46. Liesch, J. M., Sweeley, C. C., Staffeld, G. D., et al. (1982) Structure of HC-toxin, a cyclic tetrapeptide from *Helminthosporium carbonum*. Tetrahedron 38, 45–48.

47. Darkin-Rattray, S. J., Gurnett, A. M., Myers, R. W., et al. (1996) Apicidin: a novel antiprotozoal agent that inhibits parasite histone deacetylase. Proc. Natl. Acad. Sci. USA 93, 13143–13147.

48. Porter, N. J., Christianson, D. W. (2017) Binding of the microbial cyclic tetrapeptide trapoxin A to the class I histone deacetylase HDAC8. ACS Chem. Biol. 12, 2281–2286.

49. Christianson, D. W. (2026) Mechanism-based inhibition of zinc-dependent histone deacetylases. *RSC Chem*. Biol. 7, 1444–1455.

50. Ueda, H., Nakajima, H., Hori, Y., et al. (1994) FR901228, a novel antitumor bicyclic depsipeptide produced by *Chromobacterium violaceum* No. 968. J. Antibiot. 47, 301–310.

51. Furumai, R., Matsuyama, A., Kobashi, N., et al. (2002) FK228 (depsipeptide) as a natural prodrug that inhibits class I histone deacetylases. Cancer Res. 62, 4916–4921.

52. Taori, K., Paul, V. J., Luesch, H. (2008) Structure and activity of largazole, a potent antiproliferative agent from the Floridian marine cyanobacterium *Symploca* sp. J. Am. Chem. Soc. 130, 1806–1807.

53. Cole, K. E., Dowling, D. P., Boone, M. A., et al. (2011) Structural basis of the antiproliferative activity of Largazole, a depsipeptide inhibitor of the histone deacetylases. J. Am. Chem. Soc. 133, 12474–12477.

54. Hill, T. A., Shepherd, N. E., Diness, F., Fairlie, D. P. (2014) Constraining cyclic peptides to mimic protein structure motifs. Angew. Chem., Int. Ed. 53, 13020–13041.

55. Gavenonis, J., Sheneman, B. A., Siegert, T. R., et al. (2014) Comprehensive analysis of loops at protein-protein interfaces for macrocycle design. Nat. Chem. Biol. 10, 716–722.

56. Hosseinzadeh, P., Watson, P. R., Craven, T. W., et al. (2021) Anchor extension: a structure-guided approach to design cyclic peptides targeting enzyme sites. Nat. Commun. 12, 3384.

57. Watson, P. R., Gupta, S., Hosseinzadeh, P., Brown, B. P., Baker, D., Christianson, D. W. (2023) Macrocyclic octapeptide binding and inferences on protein substrate binding to histone deacetylase 6. ACS Chem. Biol. 18, 959–968.

58. Osko, J., Christianson, D. W. (2019) Methods for the expression, purification, and crystallization of histone deacetylase 6-inhibitor complexes. Methods Enzymol. 626, 447–474.

59. Kabsch, W. (2010) XDS. Acta Cryst. D66, 125–132.

60. Evans, P. (2006) Scaling and assessment of data quality. Acta Cryst. D62, 72–82.

61. Evans, P., Murshudov, G. (2013) How good are my data and what is the resolution? Acta Cryst. D69, 1204–1214.

62. Vonrhein, C., Flensburg, C., Keller, P., et al. (2011) Data processing and analysis with the autoPROC toolbox. Acta Cryst. D67, 293–302.

63. Vonrhein, C., Tickle, I., Flensburg, C., et al. (2018) Advances in automated data analysis and processing within autoPROC, combined with improved characterisation, mitigation and visualisation of the anisotropy of diffraction limits using STARANISO. Acta Cryst. A74, 360–360.

64. McCoy, A., Grosse-Kunstleve, R., Adams, P., et al. (2007) Phaser crystallographic software. J. Appl. Cryst. 40, 658–674.

65. Emsley, P., Lohkamp, B., Scott, W. G., Cowtan, K. (2010) Features and development of Coot. Acta Cryst. D66, 486–501.

66. Liebschner, D., Afonine, P., Baker, M., et al. (2019) Macromolecular structure determination using X-rays, neutrons and electrons: recent developments in Phenix. Acta Cryst. D75, 861–877.

67. Afonine, P., Grosse-Kunstleve, R., Echols, N., et al. (2012) Towards automated crystallographic structure refinement with phenix.refine. Acta Cryst. D68, 352–367.

68. Williams, C., Headd, J., Moriarty, N., et al. (2018) MolProbity: More and better reference data for improved all-atom structure validation. Protein Sci. 27, 293–315.

69. Choudhary, A., Gandla, D., Krow, G. R., Raines, R. T. (2009) Nature of amide carbonyl-carbonyl interactions in proteins. J. Am. Chem. Soc. 131, 7244–7246.

70. Newberry, R. W., Raines, R. T. (2017) The *n*→π* interaction. Acc. Chem. Res. 50, 1838–1846.

71. Ramachandran, G. N., Ramakrishnan, C., Sasisekharan, V. (1963) Stereochemistry of polypeptide chain configurations. J. Mol. Biol. 7, 95–99.

72. Johnson, L. N., Phillips, D. C. (1965) Structure of some crystalline lysozyme-inhibitor complexes determined by X-ray analysis at 6 Ångstrom resolution. Nature 206, 761–763.

73. McNae, I. W., Kan, D., Kontopidis, G., et al. (2005) Studying protein-ligand interactions using protein crystallography. Crystallogr. Rev. 11, 61–71.

74. Wienen-Schmidt, B., Oebbeke, M., Ngo, K., et al. (2021) Two methods, one goal: structural differences between cocrystallization and crystal soaking to discover ligand binding poses. ChemMedChem 16, 292–300.

75. White, S. P., Cohen, C., Phillips, G. N., Jr. (1987) Structure of co-crystals of tropomyosin and troponin. Nature 325, 826–828.

76. Barone, S., Eafa, A., Bello, I., et al. (2026) Structure-based design of HDAC6 selective inhibitors targeting the L2 pocket: modulating autophagy and liquid-liquid phase separation in lung cancer. J. Med. Chem., 69, 24496–24523.

77. Dowling, C. M., Hollinshead, K. E. R., Di Grande, A., et al. (2021) Multiple screening approaches reveal HDAC6 as a novel regulator of glycolytic metabolism in triple-negative breast cancer. Sci. Adv. 7, eabc4897.

78. Rodrigues, D. A., Wang, Y., Goulart Stollmaier, J., et al. (2026) Identification of a mechanism-based binding mode for a histone deacetylase 6 inhibitor. Nat. Commun. 17, 7289.

79. Ji, X., Nielsen, A. L., Heinis, C. (2024) Cyclic peptides for drug development. Angew. Chem. Int. Ed. 63, e202308251.

80. Jing, X., Jin, K. (2020) A gold mine for drug discovery: Strategies to develop cyclic peptides into therapies. Med. Res. Rev. 40, 753–810.

81. Klapars, A., Fryszkowska, A., Galanie, S., et al. (2026) Biocatalytic cascades enable manufacture of the macrocyclic peptide enlicitide. Science 392, 643–647.

82. Salvi, S., Linciano, P., Collina, S., Rossino, G. (2026) Cyclic peptides as modulators of protein-protein interactions: a survival guide from discovery platforms to AI-driven design. Int. J. Mol. Sci. 27, 6067.

83. Macalino, S. J. Y., Gosu, V., Hong, S., Choi, S. (2015) Role of computer-aided drug design in modern drug discovery. Arch. Pharm. Res. 38, 1686–1701.

